# Behavior in mice subjected to a Token Slot-machine: effect of Unpredictable Chronic Mild Stress

**DOI:** 10.1101/2021.09.12.459992

**Authors:** Arwen Emy Sfregola, Bruno Brizard, Anne-Marie Le Guisquet, Clémence Tillet, Eulalie Lefèvre, Luigino Bruni, Catherine Belzung

## Abstract

Several studies have succeeded in teaching animals (primates, pigeons, rats, but not mice) the value of tokens by having them executing a task using a vending-machine apparatus, where in order to receive the primary reinforcement (food), the animals had to perform a specific action that allowed them to obtain the secondary reinforcement (tokens: metal balls). We tried to assess this kind of behavior in mice that had previously been trained to use some tokens, with the aim of rewarding them not with food, but with other tokens, as a result of a token economy task. We found that mice exhibit economic behavior. Further on, our research tried to investigate the effect of stress on their operant decision-making. Therefore, the mice were divided into two groups: a Control group (n=10) and a group subjected to an Unpredictable Chronic Mild Stress (UCMS) treatment (n=8). We found that chronic stress increases some aspects of sub-optimal economic activity.

**Summary statement:** We designed an original model enabling to assess behavior in mice that had previously been trained to use some tokens, with the aim of rewarding them not with food, but with other tokens. Further on, our research investigated the effect of stress on their operant decision-making.

## Introduction

Gambling is the action of betting money (or an object of value) on an unpredictable event with the aim of obtaining an outcome, like money (Arthur et al, 2016). Nowadays gambling is a rather huge industry of global impact, often legalized by governments. It is a growing sector comprising commercial products such as slot-machines, bingo, lotteries, scratch cards, online poker and so on (European Commission, 2011; H2 gambling capital, 2016; STATISTA 2020). Under the pretext of creating an instrument of leisure, gambling companies, by using advanced technologies and sophisticated marketing tools, knowingly or unknowingly, risk creating a dependency that can also lead to addiction. Indeed, addiction to gambling can become an important public health issue (Shaffer, & Korn, 2002, Bastiani, 2013), that cannot simply be ignored on economic grounds.

Classified as a “Non-Substance-Related Disorders” (American Psychiatric Association, 2013; Potenza, 2014a) the *Gambling Disorder* (DG) has a high rate of comorbidity with substance use and mood disorders, such as *Major Depression* (Blaszczynski & McConaghy, 1989; Petry et al., 2005). Epidemiological studies have investigated the role of depression and/or stress in relation with gambling and gambling addiction. Cunningham-Williams and colleagues (1998) claimed that depression might precede gambling (Cunningham-Williams et al., 1998; Dixon et al., 2019). Other epidemiological studies confirm the presence of mood disorders in subjects that gamble (Blaszczynski & McConaghy, 1989, Becoña et al., 1996, Petry et al., 2005, Hodgins et al., 2005; Morasco et al., 2006; Bristow et al., 2018); and the comorbidity between stress –related disorders and gambling (Elman et al., 2010; Bischof et al., 2013; Berrault et al., 2017; Edgerton et al., 2018): in 2017, 71.7% of those with gambling reported high stress (Ronzitti et al., 2018). Rumination and failure of analytic thinking (Belzung et al., 2015) are typical traits of major depression (MDD) and are associated to gambling disorder (Berrault et al., 2017; Potenza, 2014b) in human beings. However, unfortunately the etiology of this comorbidity is not yet clear (Buchanan et al., 2020). While animal models of stress and major depression have already been validated (see for example the Unpredictable Chronic Mild Stress (UCMS) paradigm) (Willner et al., 1987; Surget, 2009, Nollet et al., 2013; Planchez et al., 2019), further investigations require operational animal models of gambling.

The literature reports some accurate experiments with rodent gamblers. Among several operant animal models to investigate the decision-making process, the «rodent Iowa Gambling Task» (r-IGT) (Bechara et al., 1994; Van den Bos et al., 2014) exercised an important role in the latest ten years: on the one hand there are experiments with manipulation of rats (Dellu-Hagedorn et al., 2018; Spruijt et al. 2000; de Visser et al. 2011; Van den Bos et al. 2014 for a review), on the other with the use of mice (Pittaras et al., 2013; Pittaras et al., 2016;).In that paradigm, some experiments used the «Mice Gambling Task (MGT) » (Pittaras et al., 2013; Pittaras et al., 2016; Pittaras et al., 2018), in which mice receive food pellets as rewards and quinine pellets as penality. However other rodent models are utilized to investigate the risk proneness and/or the tolerance of uncertain reward. Mobini and colleagues (Mobini et al., 2000) presented an experiment to analyze the rodent impulse control: rats, initially trained in operant conditioning chambers, have to press a lever to obtain food as reward. The Probabilistic-delivery Task (PDT) (Adriani & Laviola, 2006) is a rodent paradigm that allows to investigate gambling proneness: rats had to decide between small/sure versus large/luck-linked rewards. Others (Setlow et al., 2009; Simonet al., 2007) measure the levels of impulsive choice, characterized by preference for small immediate over larger but delayed rewards. With the “Rodent Slot Machine Task” (Winstanley et al., 2011; Cocker et al., 2016) Catharine Winstanley and her colleagues investigated the so-called «Near-misses» effect in rats. The literature reports that rats and humans are sensitive to the effect of the expectancy to near winning. «Near misses» encouraging the gambler to pay-in and restart the gambling session all over again. Normally the vulnerable individuals is stimulated by a salient stimulus such as light, sounds and combination of slot-machine symbols. The illusion of “near-misses” contribute to activate the brain regions related to the reward system.

All the experiments cited below showed the possibility to translate the gambling-like behavior in rodents by the use of food as positive reinforcer. Instead, there are some examples of tests in which rats receive non food rewards as intracranial self-stimulation (Tedford et al., 2014). Timothy D. Hackenberg (Hackenberg, 2009; Hackenberg, 2018) was able through the use of tokens to model economic human behavior in animals, following the rules of primary and secondary reinforcements in the context of operant conditioning (Skinner, 1953; Kelleher, 1956; Malagodi, 1967; Bullock & Hackenberg, 2006). For the purpose of our study we will focus on these. Several studies have succeeded in teaching animals (primates, pigeons, rats) the value of tokens by having them execute a task using a vending-machine apparatus (Hackenberg, 2009), where in order to receive the primary reinforcement (food), the animals had to perform a specific action that allowed them to obtain the secondary reinforcement (tokens).

The aim of our project is 1) to assess whether mice are able to develop an association between primary and secondary reinforcement (classical operant conditioning); 2) to investigate whether the secondary reinforcement can be rewarding per se (Gambling-like task); 3) to investigate the impact of UCMS on the behaviour in a gambling task. We hypothesize: 1. the secondary reward will induce an economic behaviour in the mice; 2. this will be modified by UCMS. For this purpose, we have created the “Token Slot-machine for Mice” from an instrumented observation cage based on a PhenoTyper® (©Noldus) - a traditional experimental arena for laboratory research with rodents.

## Methods

### Animals

Eighteen male C57BL/6J mice (Centre d’Elevage Janvier, France), aged 30-45 weeks, were included in the experiment. The mice were single-housed in standard laboratory cages (42cmx28cmx18cm) (to avoid aggressiveness related to food restriction) and subjected to standard laboratory conditions (12h light: 12h dark cycle, lights on at 08:30 pm, T = 21±1°C). Water was freely available. For Gambling-like task, we divided the animals into two groups in a random way: non-stressed (control) mice (n=10) or UCMS (n=8).

### Device

The Token Slot-Machine is a system composed of many connected devices (see Figure 1): 1. “Phenotyper®” produced by Noldus© with a floor arena of 30 cm × 30 cm × 43.5 cm, with an infra-red-sensitive video camera on the top. The arena settings were modified phase by phase, according to our experimental design. In total, following elements were present in the arena: a nose-poke hole, two dispensers (a food dispenser A and a token dispenser B), a small box and a token receptacle situated inside the apparatus which allowed it to assume different functions depending on mice’s behavioral responses; 2. Ethovision software (able to do noise-tail tracking of mice, calculate and interpret the raw data); 3. Noldus© remote (useful for the learning, habituation phase and gambling session); 4. Picolo Diligent© frame grabber and encoder board (able to create MPEG-4 video files); 5. Computer; 6. Black cloth to cover the Phenotyper© during the experimental session; 7. Two dispensers produced by Noldus© with height 20.0 cm (7.9’’), base diameter 10.0 cm (3.9’’) and silo diameter 7.5 cm (3.0’’) useful to dispense the food granules with weight 20 mg, 3.2 mm × 2.5 mm (0.13” x 0.10”) as primary reinforcements, or token metal balls (with the same measures of pellet foods) as secondary reinforcements.

**Figure 1:**
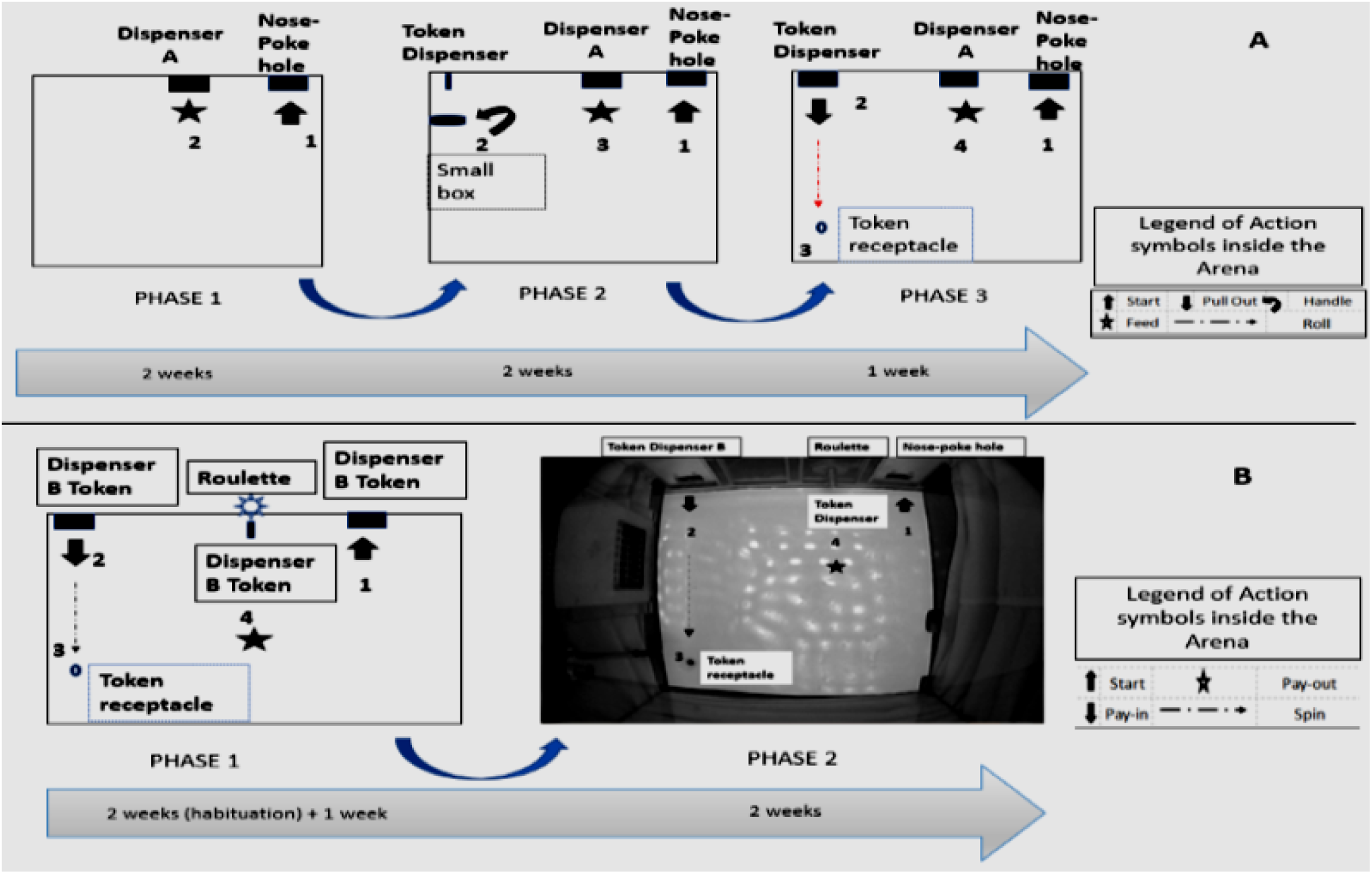
Arena settings during classical operant conditioning and gambling-like task. The numbers on the figures indicate the sequence of the different actions (1: first action of the sequence, 2: second action on the sequence, etc.) Upper panels (A): Operant conditioning: Phase 1 (left panel): mice poke their nose in the nose-poke hole in order to obtain food from Dispenser A; Phase 2 (middle panel): mice need to poke their nose in nose-poke hole in order to induce token delivery; If mice perform an action inside the small box, the food arrives in Token receptacle; Phase 3 (right panel): mice need to poke their nose in the nose-poke hole in order to induce token delivery from dispenser B (token dispenser) to token receptacle, the food arrives from food dispenser (A). Lower panels (B): Gambling-like task: Phase 1 and 2 (left panel): mice need to poke their nose in nose-poke hole in order to induce the token delivery from token dispenser B, the token falls down and enters in token receptacle; other tokens arrive in token dispenser after a roulette light; Right panel: a picture of the device

### Unpredictable Chronic Mild Stress (UCMS Treatment)

During UCMS, mice were placed in small individual cages (24cmx11cmx12cm) and exposed to a variable randomized schedule of “mild psychosocial stressors” during 10 weeks (see Table 1): baths (125 ml of water at 20°C in a cage without litter for 15 minutes), restraint stress (each mouse was kept in closed and ventilated tube of dimensions 6.5 cm length x 3.7 cm diameter), damp litter (125 ml water in each cage for a range of period of 1 or 2 hours), without litter (the litter was removed during 1 to 4 hours), social stress (each mouse was moved from its cage to one previously occupied by other individual), litter change (the litter was changed 3 times per 24 hours; the volume of each litter is 250 ml), inclined cage (45 degrees of inclinations for a duration of 1 to 2 hours) and the day-night cycle disturbances (change of the day/night cycle in four periods of 6 hours, or one to several illumination periods from 30 minutes to 2 hours during dark phase and vice-versa) (Nollet et al., 2013 ; Planchez et al., 2019). During the last five weeks behavioral tests were applied.

**Table 1:**
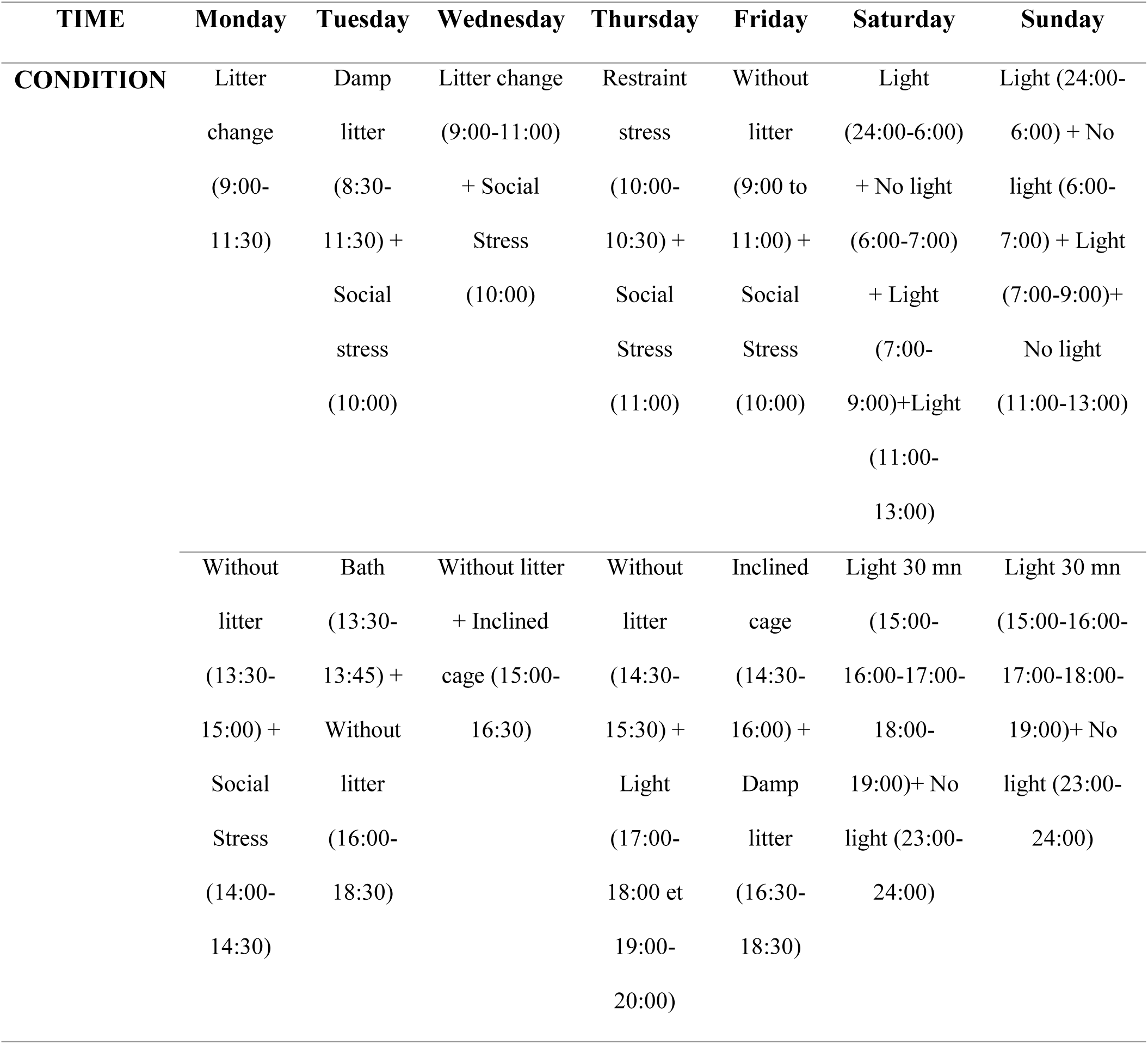
Schedule of the “Unpredictable Chronic Mild Stress”. Example of stressors schedule during the fifth week from Monday to Sunday

### Classical operant conditioning

Classical operant conditioning was undertaken in mice before the beginning of the UCMS protocol. The aim of this step was to enable mice to adopt a conditioned behavior by associating tokens with food delivery. Throughout the whole duration of this step (5 weeks in total) mice (n=18) were subjected to alimentary restriction, controlling daily that their weight did not fall under the 84% of the initial one. This step was divided into three phases and for each of them the settings of the machine were modified (see Figure 1). The variables measured in order to assess whether the mice were ready to pass from one phase to the next one were “frequency of task completion” and “latency to first performance of completion of the task”: for that the results are expressed as the time (in sec) taken by mice for the complete processing of the task. The duration of each test session for a given mouse was one hour daily for two consecutive days.

#### Phase 1

Mice learn to obtain food through step-by-step operant conditioning. Each mouse has to insert its nose in the nose-poke hole in order to obtain food from the food dispenser A.

#### Phase 2

Mice learn the association between food and token via following sequence:

a. Mice learn to insert their nose in the nose-poke hole in order to activate the token dispenser B (*Token production schedule*).
b. Mice learn to manipulate the token which has been expelled from a little tube into a box to induce activation of the food dispenser A *(Production-value schedule*). Gradually mice are exploring the token (a metal ball), and start manipulating it. The experimenter is reinforcing these behaviors with food delivery in real time.
c. Mice obtain the food from the food dispenser A *(Token exchange schedule*).

#### Phase 3

Mice are learning the value of tokens as secondary reinforcement in order to obtain food by completing following tasks:

a. Inserting their nose in hole-poke hole in order to activate the token dispenser B. (*Token production schedule*)
b. Pulling out the token using either its mouth or paw; the token is then rolling away on an inclined floor (Figure 1, Phase 3: the dashed line indicates the inclination of the plane) and fells into the token receptacle. This activates release of food from dispenser A (*Exchange production schedule*)
c. Obtaining the food from dispenser A after successfully having completed its previous behavioral response. (*Token exchange schedule*)

### Gambling-like task

This step had two main objectives: on the one hand, to enable mice to perform slot machine gambling; on the other, to check whether we find evidence of an effect of chronic stress on gambling in mice. This step was divided into 2 phases. The settings of the machine were identical for both of them, the sole difference being that we modified the ratio and schedule (with creation of pseudo-random algorithm) for receiving the token reward in Phase 2. During the phases, each reward was preceded by the alternation of a red and white light with a standard duration that modelled the roulette mechanism (see Fig.1), as a signal cue of predicted reward (Robinson, Anselme, Fischer, & Berridge, 2014). Each mouse was in the machine for one hour daily for two consecutive days. Moreover, to control the effects of UCMS that are traditionally reported in this strain, the mice were subjected to the Novelty Suppressed Feeding (NSF test) at the end of the second week (the end of “habituation”) over Phase 1. A timeline is presented in Figure 2.

**Figure 2:**
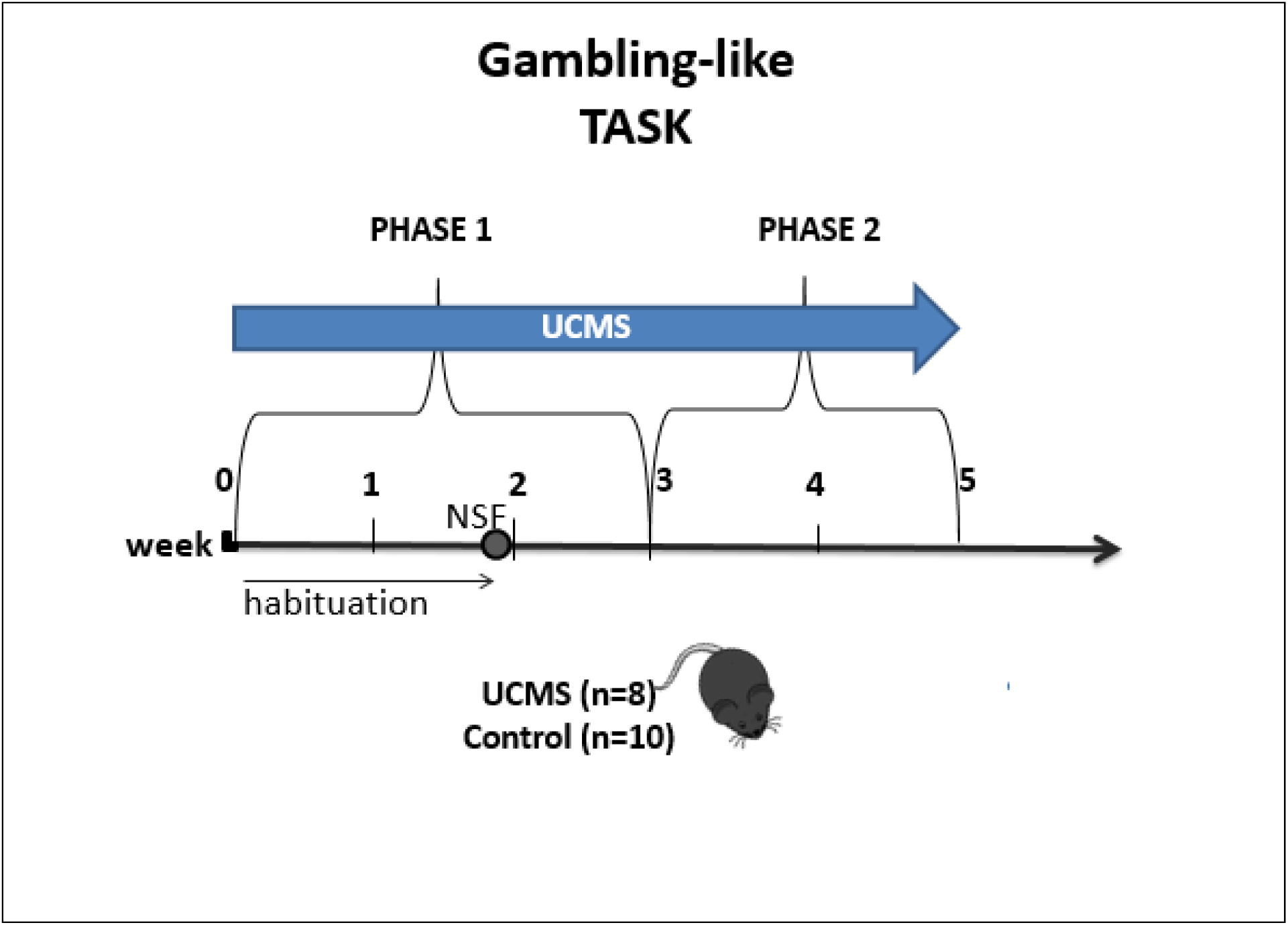
Experimental Timeline of Gambling-like task. UCMS: Unpredictable Chronic Mild Stress, NSF: Novelty Suppression of Feeding

#### Phase 1 (Habituation and economic phase)

In this phase, the machine was changed to an Economic -machine for mice. We considered the first 4 trials (over two weeks) as a preliminary phase, called “habituation” for the mice to use tokens to obtain just tokens. At the end of habituation, we assessed the behavior of the mice in the Novelty Suppressed Feeding Test. After that we continued the sessions inside the machine over 1 week. During this phase, each mouse has to insert its nose in the Nose-poke hole in order to activate the Token dispenser B; once the token has arrived automatically in dispenser B, it has to be pulled out (“pay-in”) by the mouse so that it falls into the Token receptacle in order to activate the roulette; a reward token is then expelled (“pay-out”) from the dispenser.

#### Phase 2 (Gambling-like phase)

The arena settings of the machine as well as the task the mice have to perform to obtain the token reward was the same as in Phase 1 of the Gambling task, with an important addition: the win/loss patterns through which the mice receive the tokens (pay-out) have been modified, according a particular algorithm (see Table 2). Our aim was to transform the Phenotyper from an Economic-machine to a Slot-machine for mice.

**Table 2:**
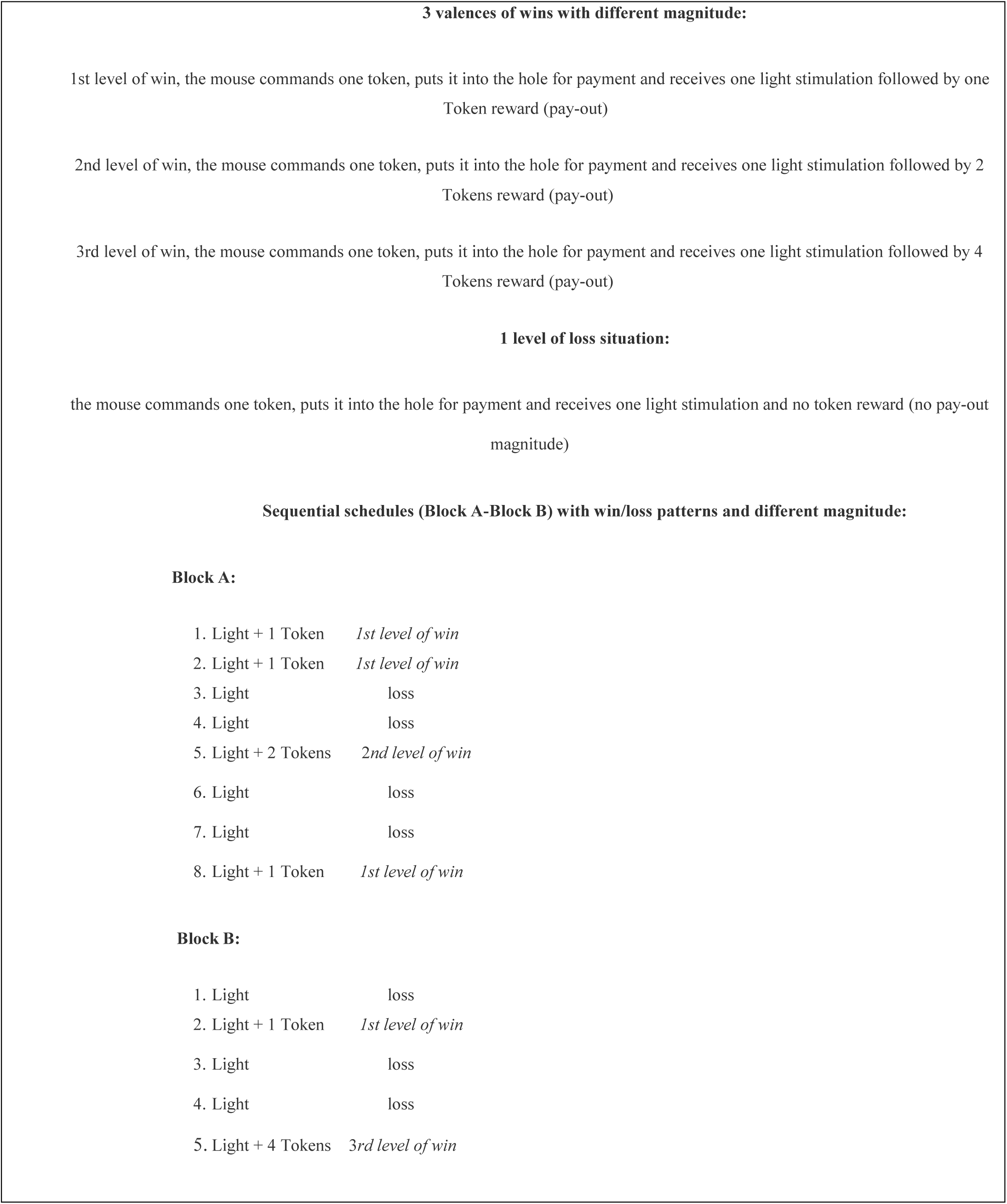
Step 2 Phase 2. Schedule of win/loss pseudo-random algorithm

In both Phases we measured the mice performance through the variable “Frequency of task completion”.

### Novelty Suppression of Feeding Test

Food was removed from the cages 12h before the test. The experimental arena consisted of a PVC box (30×30×20cm) illuminated by a red light (10 lux) and a clean sawdust as floor. At the time of testing (individual session), a single pellet of regular food was placed at the center of the box on a little white paper. The mouse was placed in a corner and confronted to a conflict between two actions: exploring the new environment and the driving to eat the pellet food within 3 minutes. We measured: latency to first smelling, frequency of smelling, latency to crunch. After the performance in the box, the mouse had to stay in its cage with the croquette for 5 minutes. In the end of the stage the quantity of food eaten was weighted.

### Statistical analysis

During classical operant conditioning, in order to assess learning of the mice, we used as an indicator the Pearson product moment correlation between the following variables: “frequency of task completion” and “latency to first performance of completion of the task”. In particular, we set as a sufficient condition and threshold of passing from one phase to the other, the registration during the last performed trial of a negative correlation between these two indicators of at least - 0.30 showing a decreasing time of “latency to first performance of completion of the task” and increasing “frequency to task completion”. For the gambling-like task, we first analyzed the performance and treatment effect in each group by applying the non-parametric Mann-Whitney test. The results are expressed as the Mean +/- SEM (Standard error of the mean). We then tested the impact of the different treatments over time on the performance of the control and UCMS group by applying the repeated-measure ANOVA test. We then conducted three different conservative F-tests for the results of Gambling-like task, Phase 2: 1) Huynh-Feldt, 2) Greenhouse-Geisser and 3) Box’s conservative F.

## Results

### Classical operant conditioning

As Phase 1 and 2 of this phase correspond to classical operant conditioning, results are not displayed here. The Pearson product moment correlation between variables: “frequency of task completion” and “latency to first performance of completion of the task” moved from statistically insignificant and slightly positive to statistically significant and strongly negative with values in the last trials of phase 1, 2 and 3 respectively: -0.56, -0.31 and -0.37 respectively. This trend confirmed that mice had gained enough comprehension and ability to handle the task before moving on to the next stage of the experiment.

We also took into consideration the last phase of classical operant conditioning (Phase 3) in order to assess whether, before starting the gambling-like step, the performance of the mice of the future groups did not differ regarding the variable “frequency of task completion”. We did not find any significant difference between the performance of the two groups of mice (control and UCMS) in trial 1 (P= 0.3954) and in trial 2 (P= 0.4222) (results not shown).

### Gambling-like task

#### Phase 1 (Habituation and economic phase, see Figure 3)

We confirmed that both groups (control and UCMS) understood the task before concluding this first phase (trial 6): all mice achieved a minimal score of 2 for “frequency of task completion”. By applying the Mann-Whitney test we do not find any significant difference between the performance of the control and UCMS group (trial 1: P= 0.4702); trial 2: P= 0.2042; trial 3: P= 0.4166; trial 4: P= 0.2964; trial 5: P= 0.1866; trial 6: P= 0.2225).

**Figure 3:**
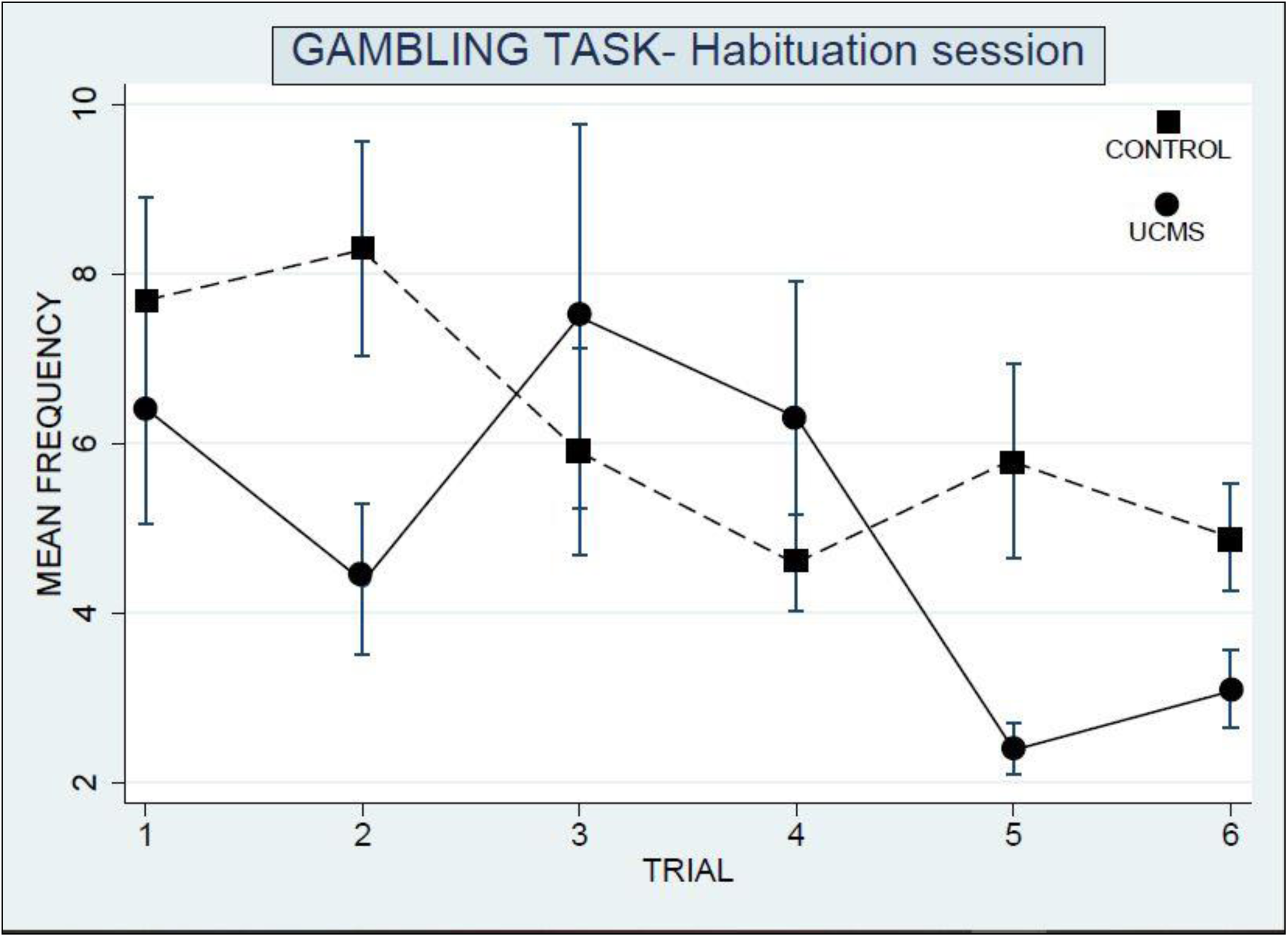
Gambling-like task, Habituation and Economic Phase. Frequency of task completion of control and UCMS groups: no significant differences are found. Mice from both groups continued to use secondary reinforcement to obtain the tokens as reward. Data represent Mean+/-SEM.

We verified also the progression over time (repeated trials) using repeated-measure ANOVA test: again, we did not find, for both of the two groups, statistically significant difference between the frequency of the task completion in the subsequent trials with respect to the first one (P= 0.4674).

#### Novelty Suppressed Feeding (NSF) test (Figure 4)

At the end of the habituation phase, we tested the mice’s anxiety behavior by applying the Novelty Suppressed Feeding test. We found a statistically significant difference between the two groups with regard to the Frequency of smelling, the performance of the UCMS mice being lower than the one of the control group (P= 0.009). Other variables were not different among groups (Latency to eat: P=0.960; Latency to smell: P=0.213; Food consumption: P=0.563). Such result allowed us to confirm that the UCMS treatment had significant impact on mice performance.

**Figure 4:**
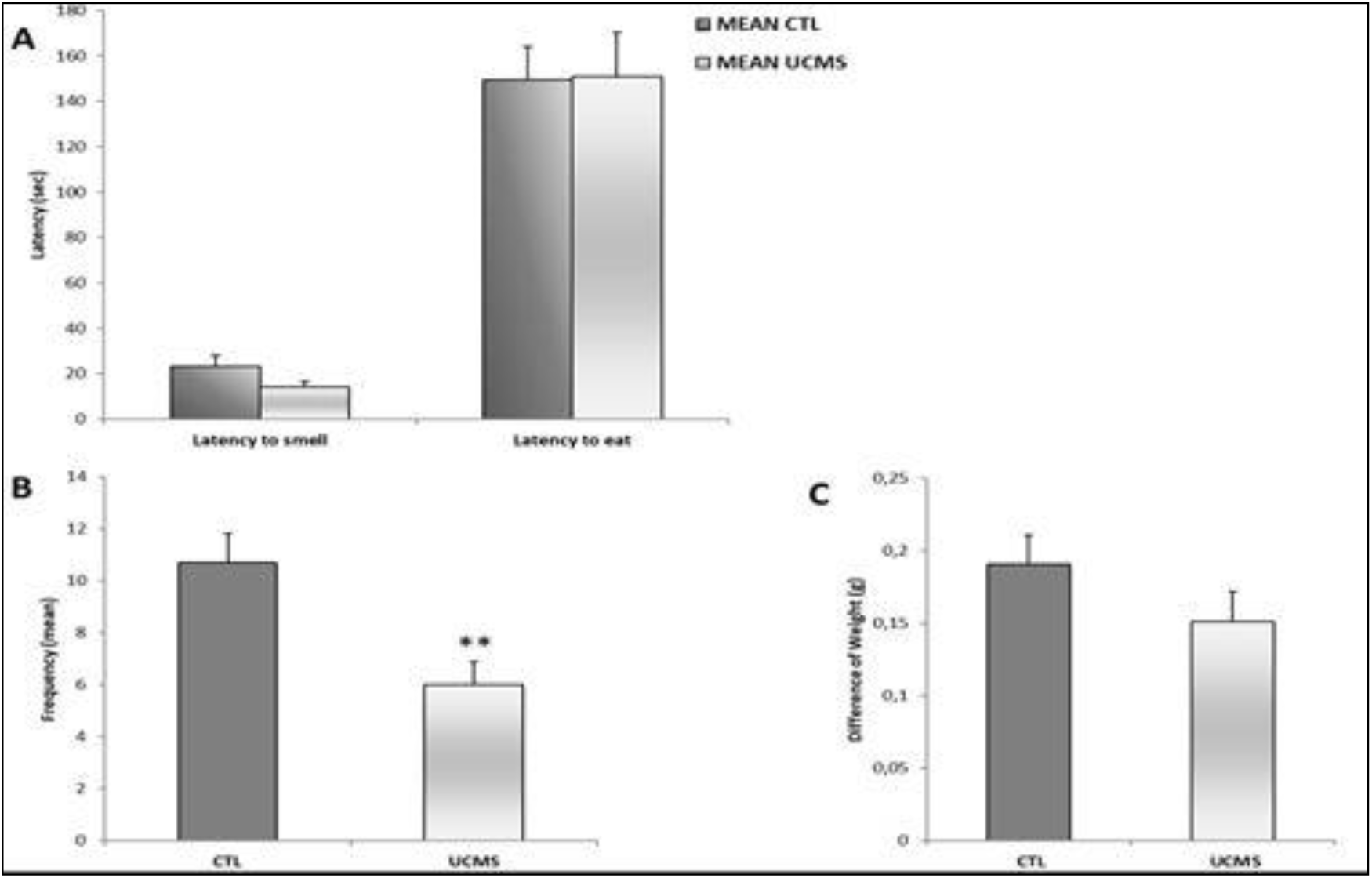
Effect of Unpredictable Chronic Mild Stress (UCMS) in the Novelty suppression of Feeding test. (A) Latency to smell and to eat). (B) Frequency to smell food before eating (C) Food consumption within 5 minutes. Results are Mean +/-SEM **P=0.009 (Mann-Whitney).

#### Phase 2 (Gambling-like phase, see Figure 5 and 6)

We found a significant difference between the performance of the two groups in the first trial (control and UCMS) with a P = 0.0066 (Means+/- SEM). No statistically significant difference between the two groups was found in the subsequent trials.

**Figure 5:**
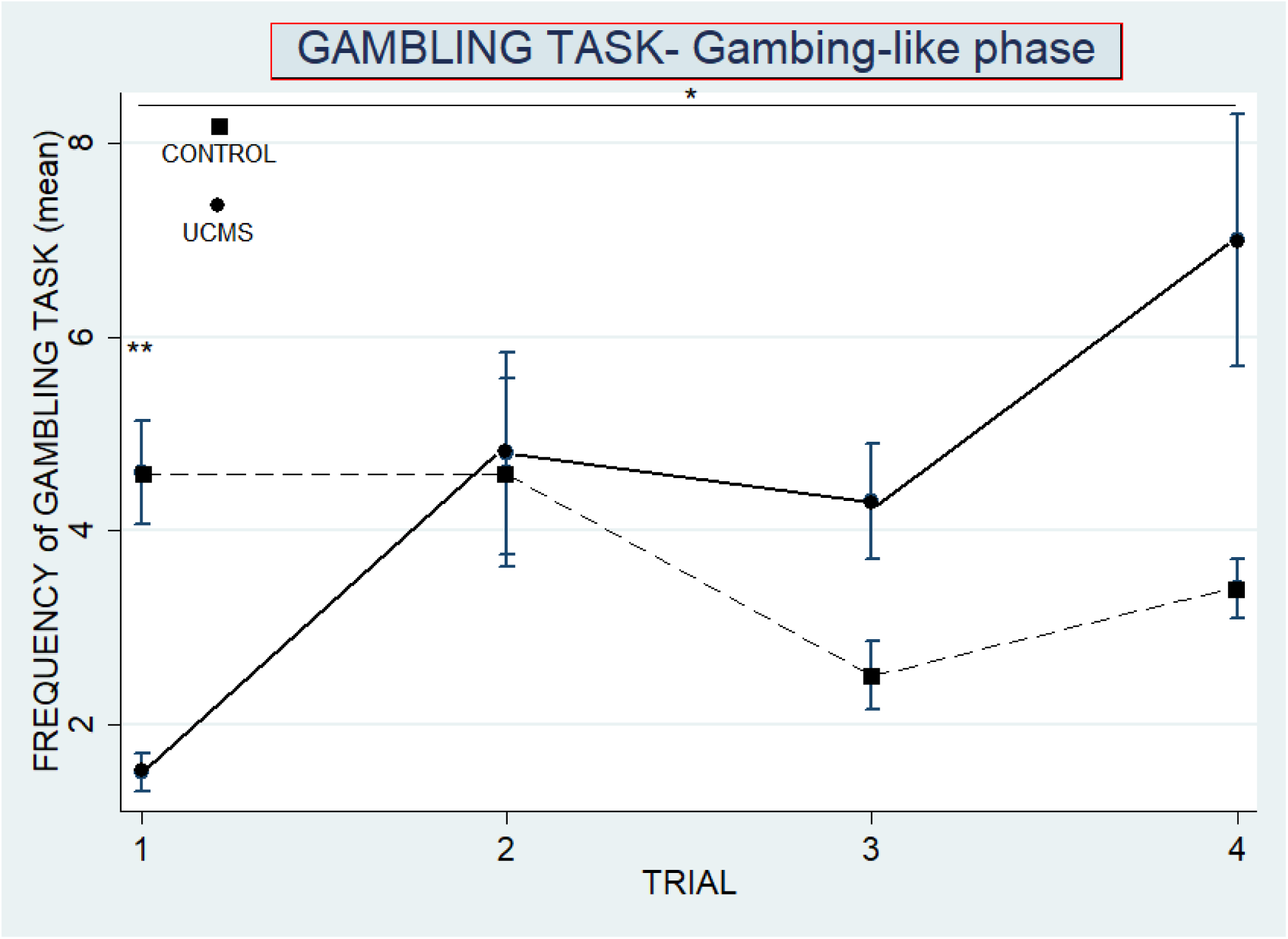
Frequency of gambling (Gambling-like task, Gambling-like phase) Significant difference was registered during the first trial (**P = 0.0066, Mann-Whitney test). Results are Means+/- SEM; UCMS: Unpredictable Chronic Mild Stress

**Figure 6:**
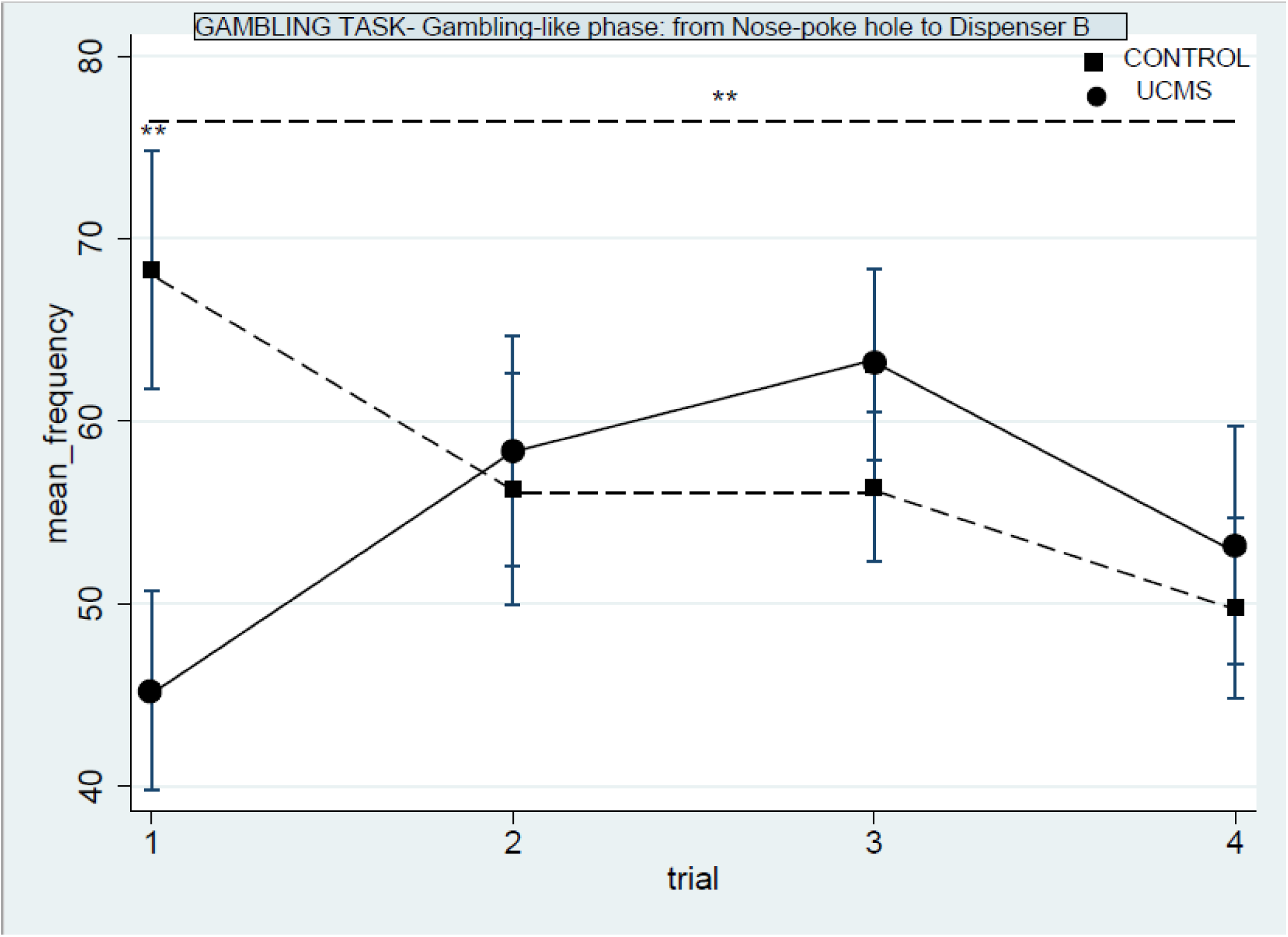
Frequency of movement from Nose-poke hole to Token dispenser B, proneness to gamble (gambling-like step, gambling phase). Initial performance **P= 0.0099. Treatment-by-time interaction: **P=0.0012. Results represent Mean+/-SEM. UCMS: Unpredictable Chronic Mild Stress)

By applying the ANOVA for repeated measures test, we detected that the treatment-by-time interaction was significant: F (3, 48) = 3.42 P =0.0244, a result confirmed also by the additional two different conservative F-tests: 1) Huynh-Feldt (P= 0.0330), 2) Greenhouse-Geisser (P= 0.0436).

We have decided to measure the mean frequency of movement from Nose-poke hole to Token dispenser B that the two groups made for each trial over the time, in order to examine the gambling proneness of the mice. We found a significant difference between the initial performance of the two groups (control and UCMS) with a P = 0.0099. No statistically significant difference between the two groups was found in the subsequent trials.

By applying the ANOVA for repeated measures test we detected that the treatment-by-time interaction was significant: F (3, 48) = 6.20 and P=0.0012.

## Discussion

We designed a mouse model enabling to assess gambling-like behavior in a context of operant conditioning under a pseudo-random algorithm (see Table 2). Over time we obtained response acquisition and maintenance of the gambling-like behavior by the use of the token as secondary reinforcement (Skinner, 1956, Malagodi, 1967; Hackenberg, 2009).

### A chained economic-system of reinforcement

Starting from this point, we can examine our setup as a chained system composed of different schedules of reinforcement (Kelleher, 1956; Jwaideh, 1973; Hackenberg, 2009): (i) start: the mouse response produce the fall of the token corresponding to the motivation of the mice to activate an economic process performed by the machine; (ii) pay-in: the possibility to exchange the token is made available in order to spin later on; (iii) pay-out: the token is being expelled as a reward. Regarding the latter, it should be noticed that once the token has been delivered (pay-out), it vanishes inside the token receptacle in order to activate again the chained economic system of reinforcement. In this sense, the reward pay-out is not just a “cold” reward, but assumes the function of an incentive salience event (Berridge & Robinson, 1998), because it can induce in mice a further reward-seeking behavior to trigger again the economic cycle. The disappearance of the final reward becomes attractive as a ‘motivational magnet’ (Olney et al., 2018) for mice.

### Sub-optimal tokens economy for mice: Gambling-like task, phase 1

According to the results for the gambling-like task phase 1, there is no difference between the performance of the Control and UCMS group in this stage of the experiment: both groups went through the learning phase of habituation, after which we observe the decline in their interest to activate the machine. In any case, mice have learned that to obtain a final reward it is necessary to perform an economic action (an effort) which finally allows them to pay-in and receive the token.

Our experiment is a pilot study regarding the possibility that the mice use the token as a secondary reinforcement. Furthermore, such economic transaction is a sub-optimal option for mice, where the effort (pay-in) equals the reward (pay-out) against to every natural action of interest (Paglieri et al., 2014; Kalenscher, & van Wingerden, 2011). Such a finding opens the searching towards comparative perspective between humans and animals, in which the behavioral psychologist can analyze the economic behavior by using an adapted animal lab vending machine (Skinner, 1956). What Skinner referred to as an example of economic control when analyzing the power of the market (and its products) for humans, goes for rodents, too.

### The effect of Unpredictable Chronic Mild Stress treatment

According to the results obtained from the Novelty Suppression of Feeding test, the UCMS mice displayed significantly lower frequency to smell the pellet food in the center of the open-field compared to the control group. The finding reveals that UCMS influenced the behavior of mice under three ways, that highlight the response to an uncertain environment : (i) greater risk-proneness (as eating a food-pellet with less precautions reflected in the reduced frequency of smelling the pellet) (Barfield et al., 2013) (ii) the repetition of schemata learned during classical conditioning, in which the animals associated the exploration of a new environment with obtaining food as primary reward (Hamilton & Brigman, 2015; Bessa et al., 2009) (iii) as a way of countering the adverse effects of uncertainty, with the motivation of a “need to know” and extract exploitable information from an unpredictable environment (Robinson, 2019).

### Slot-machine for mice as a multiple-schedule arrangement of reinforcement: Gambling-like task, Phase 2

As already noted, during Gambling-like task, Phase 2 the mice activated the multiplied-schedules of reinforcement referred to as a “slot-machine for mice” through their behavioral responses, with an important additional element: the rewarding by an algorithm based on win/loss patterns perceived as pseudo-random by mice. Before examining the presence or absence of significant effects of stress over time, we want to focus on the win/loss patterns arranged for the slot-machine project and their effects in the first and second trial of the gambling-like phase. We observed a difference between the two groups in the first trial which is in accordance with our expectation: on the market the most attractive slot-machines are the ones composed by a particular distribution of wins of reinforcement (the effect of early wins and the unforced trials is considered attractive) (Haw, 2008). Our control mice continued to play in this manner for the second trial as well, but not afterwards: in our case the novelty of the game was attractive only at the beginning. The UCMS group responded with lower mean frequency at the first trial, before changing their behavioral responses in the second trial. In the second trial, the two groups performed equally.

### Performance and the motivation of the UCMS and control mice over time in a gambling-like task, Phase 2: for what kind of reward?

There is no difference between the two groups over time in regard to the mean frequency of the complete gambling-like task, Phase 2. Nevertheless, UCMS mice exhibited increased behavioral responses trial by trial. This finding could reveal a possible attraction to a system of reinforcement under chronic stress condition, in which they are not sure about the future wins. This is known as the phenomenon of “the attractiveness of reward uncertainty”, as explained by the compensatory hypothesis (Anselme & Robinson, 2013). This hypothesis states that when animals are in a situation of uncertainty as physiological deprivations, psychosocial deprivations, their motivational resources are brought out in order to respond to this kind of situation by doing something rather than nothing, even if this behavior is sub-optimal.

We have also checked the proneness of mice to activate the start schedule, and observed that the Control mice lost their preference to gamble over time while the UCMS mice only decreased their movements from Nose-poke hole to Token dispenser B during the last week, probably because they became more able to pay-in order to gamble.

## Conclusion

Gambling can be viewed as the uncertain and pseudo-random schedule of reward that the animals, in depressive-like state, are able to activate over time. We do not, however, know whether this sub-optimal economic behavior is simply a compensative response or whether they do it for pleasure as well. As we asserted above, in accordance with the incentive salience hypothesis, probably the pay-out is not a neutral reward but is related instead to the motivational components and the role of dopamine as an incentive salience event, especially when the ratio of reinforcement is subjected under the pseudo-random algorithm (Winstanley et al., 2011; Robinson et al., 2015). In order to confirm such hypothesis, we should investigate the neurobiological underpinnings of mice behavior.

## Acknowledgements

We wish to express our appreciation to the following for their helpful comments, and provocative and illuminating discussions about our project: Elisabetta Benedetto, Stefania Cosci, Alessandro Giuliani, Benedetto Gui, Marc Legrand, Angeline Maillard, Antonello Maruotti, Erica Mastrociani, Bill Neu, Paul O’Hara, Vittorio Pelligra, Barbara Planchez, Fabio Ritossa, Alexandre Surget, Romain Troubat, Agnes Vermogen.

We are indebted to Marta Pancheva for comments on the use of statistics, for statistical assistance with the STATA Software, and for her comments and suggestions on the manuscript.

We are also very grateful to Alessandro Svetina, for the technical assistance in the layout of the figures.

## References

Adriani, W., & Laviola, G. (2006). Delay aversion but preference for large and rare rewards in two choice tasks: implications for the measurement of self-control parameters. BMC Neuroscience 7, 52. http://doi.org/10.1186/1471-2202-7-52

American Psychiatric Association (2013). Diagnostic and Statistical Manual of Mental Disorder, (5th ed.).

Anselme., P. & Robinson, M. J. F. (2013). What motivates gambling behavior? Insight into dopamine’s role. Frontiers in Behavioral Neuroscience, 7, 182. http://doi.org/10.3389/fnbeh.2013.00182

Anselme P, Güntürkün O. (2018) How foraging works: uncertainty magnifies food-seeking motivation [published online ahead of print, 2018 Mar 8]. Behav Brain Sci.;1‐106. http://doi./org/10.1017/S0140525X18000948

Arthur, J. N. Williams, R. J., & Delfabbro, P. H. (2016). The conceptual and empirical relationship between gambling, investing, and speculation. Journal of Behavioral Addictions, 5, 580–591. http://doi.org/10.1556/2006.5.2016.084

Barfield, E.T., Moser, V.A., Hand, A., & Grisel, J.E. (2013). β-endorphin modulates the effect of stress on novelty-suppressed feeding. Front. Behav. Neurosci., 7(19), 1–7. http://doi.org/10.3389/fnbeh.2013.00019

Bastiani, L., Gori, M., Colasante, E., Siciliano, V., Capitanucci, D., Jarre, P., Molinaro, S. (2013). Complex Factors and Behaviors in the Gambling Population of Italy, J Gambl Stud. 29, 1–13. http://doi.org/10.1007/s10899-011-9283-8

Bechara, A., Damasio, A. R., Damasio, H, & Anderson, S. W. (1994). Insensitivity to future consequences following damage to human prefrontal cortex. Cognition 50 7–15. http://doi.org/10.1016/00100277(94)90018-3

Becoña, E., Del Carmen Lorenzo, M. & Fuentes, M. J. (1996). Pathological gambling and depression. Psychological Reports 78, 635–640. http://doi.org/10.2466/pr0.1996.78.2.635

Belzung, C., Willner, P. & Philippot, P. (2015). Depression: from psychopathology to pathophysiology, Current Opinion in Neurobiology, 30, 24–30. http://doi.org/10.1016/j.conb.2014.08.013

Berrault, S., Bonnaire, C., & Herrmann F. (2017). Anxiety, Depression and emotion regulation among regular online poker players, J Gambl Stud. 33, 1039–1050. http://doi.org/10.1007/s10899-017-9669-3

Berridge, K. C., & Robinson, T. E. (2016). Liking, wanting and the incentive-sensitization theory of addiction. The American psychologist 71, 670–679. http://doi.org/10.1037/amp0000059

Berridge, K.C., & Robinson, T.E. (1998). What is the role of dopamine in reward: hedonic impact, reward learning, or incentive salience?, Brain Res Brain Res Rev. 28, 309–69.

Bessa, J. M., Mesquita, A. R., Oliveira, M., Pêgo, J. M., Cerqueira, J. J., Palha, J. A., Almeida O. F. X., Sousa, N. (2009) Trans-Dimensional Approach to the Behavioral Aspects of Depression. Frontiers in Behavioral Neuroscience 3, 1. http://doi.org/10.3389/neuro.08.001.2009

Bischof, A., Meyer, C., Bischof, G., Kastirke, N., John, U., Rumpf, H. J. (2013). Comorbid AxisI-disorders among subjects with pathological, problem, or at-risk gambling recruited from the general population in Germany: results of the PAGE study. Psychiatry Res. 210, 1065–70. http://doi.org/10.1016/j.psychres.2013.07.026

Blaszczynski,, A, & McConaghy, N. (1989). Anxiety and/or depression in the pathogenesis of addictive gambling, Int J Addict. 24, 337–350.

Bristow, L. A., Bilevicius, E., Stewart, S.H. Goldstein A.L. & Keough M.T. (2018), Solitary gambling mediates the risk pathway from anxiety sensitivity to excessive gambling: evidence from a longitudinal ecological momentary assessment study, Psychol Addict Behav, 32, pp. 689–696

Buchanan, T. W., McMullin, S. D., Baxley, C. & Weinstock, J. (2020) Stress and gambling, Current Opinion in Behavioral Sciences, 31, 8–12, http://doi.org/10.1016/j.cobeha.2019.09.004

Bullock, E. C., & Hackenberg, T. D. (2006). Second-Order Schedules of Token Reinforcement with Pigeons: Implications for Unit Price, J Exp Anal Behav. 85, 95–106.

Cocker, P.J., Vonder Haar, C., & Winstanley, C.A. (2016). Elucidating the role of D4 receptors in mediating attributions of salience to incentive stimuli on Pavlovian conditioned approach and conditioned reinforcement paradigms, Beh. Brain Res. 312, 55-63. doi: http://doi.org/10.1016/j.bbr.2016.06.007

Coriale, G., Ceccanti, M., De Filippis, S., Falletta Caravasso, C., & De Persis, S. (2015). Disturbo da gioco d’azzardo: epidemiologia, diagnosi, modelli interpretativi e trattamento (Gambling Disorder: epidemiology, diagnosis, interpretative models and intervention), Riv Psichiatr 50 216–227.

Cunningham-Williams, R. M., Cottler, L.B., Compton, W. M., & Spitznagel, E. L. (1998). Taking chances: problem gamblers and mental health disorders-results from the St. Louis Epidemiologic Catchment Area Study. American Journal of Public Health 88, 1093–1096. http://doi.org/10.2105/AJPH.88.7.1093

Dellu-Hagedorn F, Rivalan M, Fitoussi A, De Deurwaerdère P. (2018) Inter-individual differences in the impulsive/compulsive dimension: deciphering related dopaminergic and serotonergic metabolisms at rest. Philos Trans R Soc Lond B Biol Sci. 373(1744):20170154. doi:10.1098/rstb.2017.0154

De Visser, L., Homberg, J. R., Mitsogiannis, M., Zeeb, F. D., Rivalan, M., Fitoussi, A., & Dellu-Hagedorn, F. (2011). Rodent Versions of the Iowa Gambling Task: Opportunities and Challenges for the Understanding of Decision-Making, Frontiers in Neuroscience, 5, 109. http://doi.org/10.3389/fnins.2011.00109

Dixon, M.J., Gutierrez, J., Stange, M., Larche, C.J., Graydon, C., Vintan, S. & Kruger, T. B., (2019), Mindfulness problems and depression symptoms in everyday life predict dark flow during slots play: implications for gambling as a form of escape, Psychol Addict Behav, 33, pp. 81–90

Edgerton, J.D., Keough M.T. & Roberts L.W. (2018) Co-development of problem gambling and depression symptoms in emerging adults: a parallel-process latent class growth model, J Gambl Stud, 34, pp. 949-

Elman, I., Tschibelu, E. & Borsook, D. (2010). Psychosocial stress and its relationship to gambling urges in individuals with pathological gambling. The American journal on addictions19 4, 332–9. http://doi.org/10.1111/j.1521-0391.2010.00055.x

European Commission (2011), Green Paper, On on-line gambling in the Internal Market. http://eurlex.europa.eu/legalcontent/EN/TXT/PDF/?uri=CELEX:52011DC0128&from=EN/

H2 gambling capital 2016, retrieved from http://h2gc.com/news/reports/h2-features-as-ecomomists-graph-of-the-day-for-the-third-time-to-coincide-with-the-opening-of-ice/2016

Hackenberg, T. D. (2009). Token Reinforcement: a review and analysis, Journal of the experimental analysis of behavior, 2, 257–286. doi: 10.1901/jeab.2009.91-257

Hackenberg TD. Token reinforcement: Translational research and application. J Appl Behav Anal. 2018;51(2):393‐435. doi:10.1002/jaba.439

Hamilton, D. A.; & Brigman, J. L. (2015). Behavioral flexibility in rats and mice: Contributions of distinct frontocortical regions. Genes, Brain, and Behavior, 14, 4–21. http://doi.org/10.1111/gbb.12191.

Haw, J. (2008). Random-ratio schedules of reinforcement: The role of early wins and unreinforced trials, Journal of Gambling Issues 21, 56–67. http://doi.org/10.4309/jgi.2008.21.6

Hodgins, D. C., Peden, N., & Cassidy, E. (2005). The Association between comorbidity and outcome in pathological gambling: a prospective follow-up of recent quitters. J Gambl Stud., 2, 255–71. http://doi.org/10.1007/s10899-005-3099-3

Jwaideh, A. R. (1973). Responding under chained and tandem fixed-ratio schedules. Journal of the Experimental Analysis of Behavior 19(2), 259–267. http://doi.org/10.1901/jeab.1973.19-259

Kalenscher, T., & van Wingerden, M. (2011). Why we should use animals to study economic decision making - a perspective. Frontiers in neuroscience, 5, 82. http://doi.org/10.3389/fnins.2011.00082

Kelleher, R. T. (1956). Discrimination learning as a function of reversal and nonreversal shifts. Journal of Experimental Psychology 51, 379–384. http://doi.org/10.1037/h0047413

Malagodi, E. F. (1967). Fixed-ratio schedules of token reinforcement, Psychon. Sci. 8: 469. http://doi/10.3758/BF03331702

Mobini, S., Chiang, T. J., Al-Ruwaitea, A. S. A., Ho, M. Y., Bradshaw, C. M. & Szabadi, E. (2000) Effect of central 5-hydroxytryptamine depletion on inter-temporal choice: a quantitative analysis. Psychopharmacology 149, 313–318. http://doi.org/10.1007/s002130000385

Morasco, B. J Vom Eigen, K. A, & Petry, N. (2006). Severity of gambling is associated with physical and emotional health in urban primary care patients. General hospital psychiatry 28, 94–100. http://doi.org/10.1016/j.genhosppsych.2005.09.004

Nollet, M., Le Guisquet, A. M. & Belzung, C. (2013), Models of depression: unpredictable chronic mild stress in mice, Curr Protoc Pharmacol., 61 (1), 5.65.1-5.65.17. http://doi.org/10.1002/0471141755.ph0565s61

Olney, J. J., Warlow, S. M., Naffziger, E. E., & Berridge, K. C. (2018). Current perspectives on incentive salience and applications to clinical disorders. Current opinion in behavioral sciences, 22, 59–69. https://doi.org/10.1016/j.cobeha.2018.01.007

Paglieri, F., Addessi, E., De Petrillo, F., Laviola, G., Mirolli, M., Parisi, D., Petrosino, G., Ventricelli, M., Zoratto, F., & Adriani, W. (2014). Nonhuman gamblers: lessons from rodents, primates, and robots. Front Behav Neurosci. 8: 33. http://doi.org/10.3389/fnbeh.2014.00033

Petry, N. M., Stinson, F. S., Grant, B. F. (2005). Comorbidity of DSM-IV Pathological Gambling and Other Psychiatric Disorders: Results From the National Epidemiologic Survey on Alcohol and Related Conditions, The Journal of Clinical Psychiatry 66, 564–574.

Pittaras, E., Cressant, A., Serreau, P., Bruijel, J., Dellu-Hagedorn, F., Callebert, J., Rabat, A., & Granon, S. (2013). Mice Gamble for Food: Individual Differences in Risky Choices and Prefrontal Cortex Serotonin. J Addict Res Ther S4: 011. http://doi.org/10.4172/2155-6105.S4-011

Pittaras, E., Callebert, J., Chennaoui, M., Rabat, A., & Granon, S. (2016). Individual behavioral and neurochemical markers of unadapted decision-making processes in healthy inbred mice. Brain structure & function, 221(9), 4615–4629. http://doi.org/10.1007/s00429-016-1192-2

Pittaras, E., Callebert J, Dorey R, Chennaoui M, Granon S, Rabat A., (2018) Mouse Gambling Task reveals differential effects of acute sleep debt on decision-making and associated neurochemical changes, Sleep, 41(11), zsy168. http://doi.org/10.1093/sleep/zsy168

Planchez, B., Surget, A. & Belzung, C. (2019) Animal models of major depression: drawbacks and challenges. J Neural Transm 126, 1383–1408. https://doi.org/10.1007/s00702-019-02084-y

Potenza M. N. (2014a). Non-substance addictive behaviors in the context of DSM-5. Addictive behaviors, 39(1), 1–2. https://doi.org/10.1016/j.addbeh.2013.09.004

Potenza, M. N., (2014b). The neural bases of cognitive processes in gambling disorder, Trends Cogn Sci. 18, 429–438. http://doi.org/10.1016/j.tics.2014.03.007]

Robinson, M. J., Anselme, P., Fischer, A. M., & Berridge, K. C. (2014). Initial uncertainty in Pavlovian reward prediction persistently elevates incentive salience and extends sign-tracking to normally unattractive cues. Behavioural brain research, 266, 119–130. http://doi.org/10.1016/j.bbr.2014.03.004

Robinson M.J., Fischer A.M., Ahuja A., Lesser E.N., Maniates H. (2015). Roles of “wanting” and “liking” in motivating behavior: gambling, food, and drug addictions. In: Simpson E., Balsam P. (eds) Behavioral neuroscience of motivation. Current topics in behavioral neurosciences, vol 27, Springer, Cham.

Robinson M.J. (2019) Hoarding all of the chips: Slot machine gambling and the foraging for coins. Behav Brain Sci.;42:e50. http://doi.org/10.1017/S0140525X18001917

S. Ronzitti, S.W. Kraus, R.A. Hoff, M.N. Potenza (2018) Stress moderates the relationships between problem-gambling severity and specific psychopathologies, Psychiatry Res, 259, pp. 254–261

Shaffer, H. J. & Korn,, D.A. (2002). Gambling and related mental disorders: a public health analysis, Annu Rev Public Health 23, 171–212. http://doi.org/10.1146/annurev.publhealth.23.100901.140532

Setlow, Barry et al. (2009) Effects of chronic administration of drugs of abuse on impulsive choice (delay discounting) in animal models. Behavioural pharmacology vol. 20, 5-6: 380–9. http://doi.org/10.1097/FBP.0b013e3283305eb4

Simon, N.W., Gilbert, R. J., Mayse, J. D., Bizon, J. L & Setlow, B. (2009). Balancing Risk and Reward: A Rat Model of Risky Decision-Making. Neuropsychopharmacology : official publication of the American College of Neuropsychopharmacology, 34, 2208–2217. http://doi.org/10.1038/npp.2009.48

Simon, N. W., Mendez, I. A., & Setlow, B. (2007). Cocaine exposure causes long-term increases in impulsive choice. Behavioral Neuroscience, 121(3), 543–549. http://doi.org/10.1037/0735-7044.121.3.543

Skinner, B. F. (1953), Science and human behavior, The B. F. Skinner Foundation, https://www.bfskinner.org/product/science-and-human-behavior-pdf/

Spruijt B., (2001). A concept of welfare based on reward evaluating mechanisms in the brain: anticipatory behaviour as an indicator for the state of reward systems, Applied Animal Behaviour Science 72, 145–171.

STATISTA (2020), Market value of online gambling worldwide 2017 and 2024, https://www.statista.com/statistics/270728/market-volume-of-online-gaming-worldwide/#statisticContainer/ (accessed 2 June 2020

Surget, A., Wang, Y., Leman, S., Ibarguen-Vargas, Y., Edgar, N., Griebel, G., Belzung, C. & Sibille, E. (2009). Corticolimbic transcriptome changes are state-dependent and region-specific in a rodent model of depression and of antidepressant reversal. Neuropsychopharmacology : official publication of the American College of Neuropsychopharmacology 34; 1363–1380. http://doi.org/10.1038/npp.2008.76

Tedford, S. E., Holtz, N. A., Persons, A. L., & Napier, T. C. (2014). A new approach to assess gambling-like behavior in laboratory rats: using intracranial self-stimulation as a positive reinforcer. Frontiers in behavioral neuroscience, 8, 215. http://doi.org/10.3389/fnbeh.2014.00215

Van den Bos, R., Koot, S. & de Visser, L. (2014). A rodent version of the Iowa Gambling Task: 7 years of progress. Frontiers in Psychology 5:203. http://doi.org/10.3389/fpsyg.2014.00203

Willner, P., Towell, A., Sampson, D., Sophokleous, S. & Muscat, R. (1987). Reduction of sucrose preference by chronic unpredictable mild stress, and its restoration by a tricyclic antidepressant. Psychopharmacology (Berl) 93 358–364. http://doi.org/10.1007/BF00187257

Winstanley, C. A., Cocker, P. J & Rogers, R. D. (2011). Dopamine modulates reward expectancy during performance of a slot machine task in rats: evidence for a “near-miss” effect. Neuropsychopharmacology, 36(5), 913–925. http://doi.org/10.1038/npp.2010.230

World Health Organization, (2017). Depression and Other Common Mental Disorders: Global Health Estimates (Report). Retrieved from http://www.who.int/mental_health/management/depression/prevalence_global_health_estimates/en/

